# Domain Specific AI Segmentation of IMPDH2 Rod/Ring Structures in Mouse Embryonic Stem Cells

**DOI:** 10.1101/2024.10.16.617897

**Authors:** Samuel T.M. Ball, Meagan J. Hennessy, Yuhan Tan, Kai F. Hoettges, Neil D. Perkins, David J. Wilkinson, Michael R.H. White, Yalin Zheng, David A. Turner

## Abstract

**Background:** Inosine monophosphate dehydrogenase 2 (IMPDH2) is an enzyme that catalyses the rate limiting step of guanine nucleotides. In mouse embryonic stem cells (ESCs) IMPDH2 is held as large multi-protein complexes known as rod-ring (RR) structures that dissociate when ESCs differentiate. Manual analysis of RR structures from confocal microscopy images, although possible, is not feasible on a large scale due to the quantity of RR structures present in each field of view. To address this analysis bottleneck, we have created a fully automatic RR image classification pipeline to segment, characterise and measure feature distributions of these structures in ESCs.

**Results:** We find that this model can automatically segment images with a Dice score of over 80% for both rods and rings for in-domain images compared to expert annotation, with a slight drop to 70% for datasets out of domain. Important feature measurements derived from these segmentations show high agreement with the measurements derived from expert annotation, achieving an *R*^2^ score of over 90% for counting the number of rings and rods over the dataset.

**Conclusions:** We have established for the first time a quantitative baseline for RR distribution in pluripotent ESCs and have made a pipeline available for training to be applied to other models in which RR remain an open topic of study.

## Background

### Introduction

Multiparameter fluorescence microscopy is a powerful and versatile tool in modern biology, which allows both qualitative and quantitative measurement of biological processes in fixed or live samples [1]. Despite the rapid expanse in both the availability and technological innovation seen in the field of quantitative microscopy, adequately analysing the data generated from the microscope remains a major bottleneck [2]. Artificial Intelligence (AI) however, and notably the recent advances in the application of AI based technologies, stands as a useful yet underutilised tool that could address this issue.

One potential application for deep learning techniques can be seen in the study of Inosine Monophosphate Dehydrogenase 2 (IMPDH2), the enzyme that catalyses the rate limiting step in the synthesis of guanine nucleotides, and is a critical component of cell homeostasis and metabolism [3]. IMPDH2 can exist in two major conformational classes depending on the metabolic requirements of the cell. The first is homogeneously distributed throughout the cytoplasm, usually found in in cells with a low metabolic demand (e.g. HeLa, MCF7) [4]. However, in cells that are rapidly proliferating with a high demand for nucleotides (e.g. HEp-2, mouse embryonic stem cells), IMPDH2 forms large, macromolecular complexes resembling rod or ring (RR) structures (**Fig. 1**; and [5, 6]). Interestingly, similar rod-ring structures been identified for other metabolic enzymes such as cytidine triphosphate synthase 1 (CTPS1) [5, 7], glutamate synthase [8] and GDP-mannose pyrophosphorylase [8]. This has led to the suggestion that higher-order molecular confirmations may be important for fine-tuning enzymatic kinetics independent of transcription regulation [8, 9].

**Figure 1.**
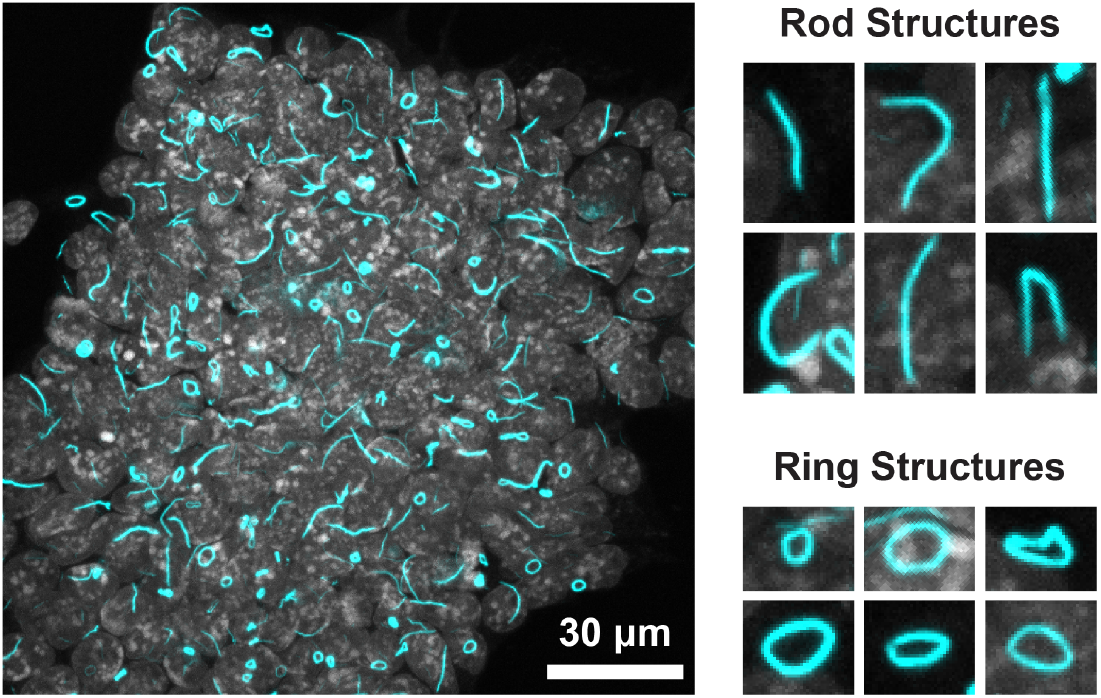
IMPDH2 rod-ring structures in pluripotent mouse embryonic stem cells. A representative maximum intensity projection of fixed mouse embryonic stem cells (left) showing DAPI (grey) which marks the nuclei, and IMPDH2 (cyan). Magnified examples of rods (top right) and rings (bottom right) are shown. The number of rods-rings in this image is in excess of 200. Scale bar indicates 30 µm.

Mouse ESCs are a useful model system to study the regulation of these structures, as the IMPDH2 conformation differs depending on their state (**Fig. 1**; and [5, 6]). When ESCs are maintained in pluripotent conditions IMPDH2 is mostly in the rod-ring conformation. During differentiation however, these structures break down and IMPDH2 becomes more homogenously cytoplasmic. As these structures have a clear morphology, single instances of rods and rings are relatively straightforward to analyse, and manual quantification similar to our previously published protocols [10, 11] or simple thresholding using, for example ImageJ/FIJI is possible [12-14]. However, within a single field of view there can be dozens to over a hundred rods and rings, rendering manual quantification impractical, especially if one is to analyse enough structures for meaningful statistical analysis.

### Software Requirements

Due to the number of these structures within each image (**Fig. 1**) and the impractical nature of current analysis methods, it is clear that any software that could tackle our analysis requirements would need to:

1. Identify rods or rings with a high degree of accuracy, consistently over multiple experimental replicates, conditions, and fluorescence intensities compared with manual identification;
2. Be automated, requiring little to no adjustments or corrections;
3. Run through datasets at least an order of magnitude quicker than manual analysis;
4. Be user-friendly, and able to export the data in a format that can be interrogated by third party graphing/analysis packages.

We therefore set out to develop a deep learning approach to establish a numerical baseline for IMPDH2 rod and ring distribution in mice. Since these rod-ring structures are present in multiple contexts (e.g. IMPDH2 [5], CTPS1 cytoophidia [5, 7], Loukoumasomes [15]), the solution we develop is expected be of use to multiple groups for the quantitative analysis of similar cellular structures.

### Related work and Background Theory

Image segmentation is a commonly investigated computer vision task, with models for object detection via bounding boxes [16], prompt based segmentation [17] and semantic segmentation [18]. The methodology for using these models is typically split into two cases: in the first; pretrained models are used directly on new datasets to detect objects or segment images. In the second, these model architectures are partially, or fully retrained on the new data to achieve greater results.

Commonly used applied models such as YOLO [16] or SegFormer [18] come with labels for general use (e.g. dog, person, train), but are inappropriate for rod ring segmentation as there is no “rod” or “ring” output in the pretrained model outputs. The large size of these models, combined with the relatively small size of some microscopy imaging datasets typically result in these larger architectures being outperformed by simpler models at a smaller scale [19]. Other models, such as Segment Anything [17] or Cellpose [20] take a label-free approach to segmentation, using user prompting to infer structure in the image. While these models cannot achieve end-to-end segmentation like those above, they are still used to increase the speed of segmentation and annotation for many biomedical imaging tasks.

Training bespoke, application specific models allow for specific considerations to be made for the type of data being analysed, such as preprocessing or model design. By training a model on a specific dataset, higher performance is usually achieved compared to generalised models due to a smaller domain space; however, performance out-of-domain (i.e. on data sufficiently dissimilar to the training data) is typically much worse.

### Implementation

#### AI Methods

##### Training

For training the microscopy AI models, we used 5 UNet models [21] in an ensemble model (**Fig. 2A**). Each input image is scaled to 512×512 pixels, and the output logits of the UNet models are averaged to give an aggregate logit map of the output. For downstream binary classification tasks, the output was thresholded at a value of 0.2; chosen due to the ensemble model’s propensity to over-classify negative classes (i.e. empty space) in the image. The dataset totalled 287 training images, each with an accompanying annotation mask for rings and rods (**Fig. 2B**). This was split into 5 disjoint folds, where each model was trained on 4 of these folds and tested on the last one, with each model testing on a different fold. This ensured complete testing coverage of the entire dataset, and repeated training to ensure reliability of the protocol.

**Figure 2.**
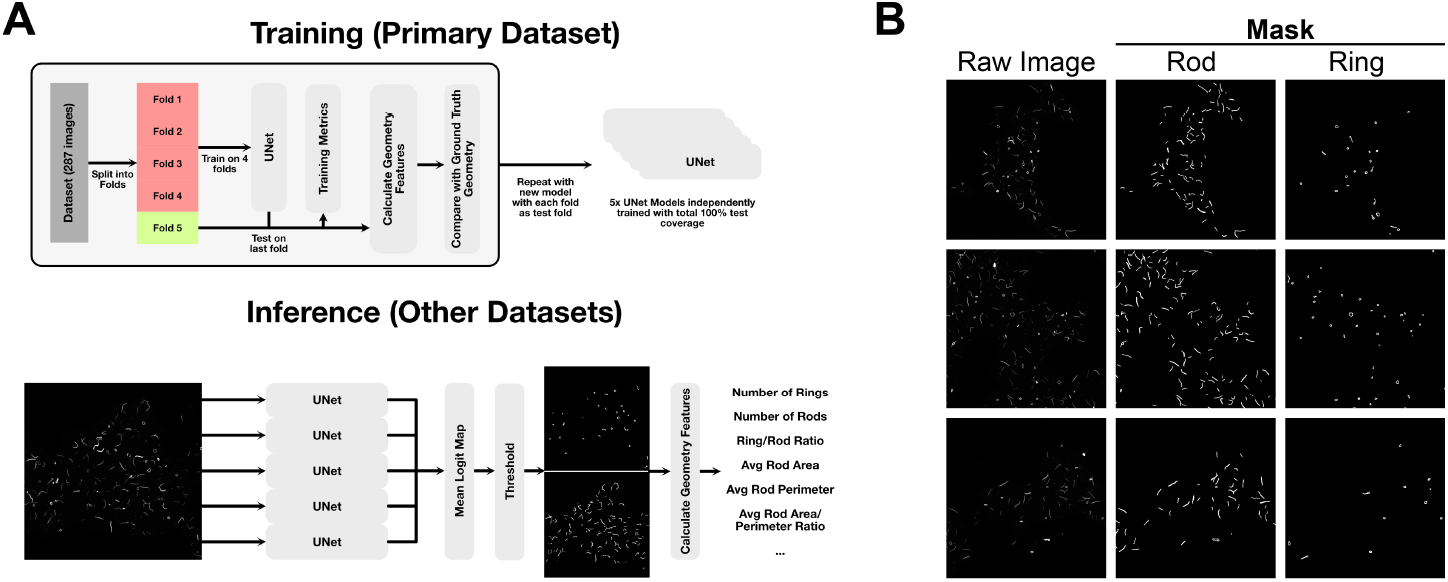
Training and inference protocol, and example training labels for rod ring segmentation. (**A**) Training and inference protocol used in this work. The training datasets were split into 5 disjoint sets, with a UNet model trained on each combination of 4 sets, testing on the final test set each time. After the training and testing on the primary dataset is complete, this gives us 5 UNet models that are used in ensemble for inference, with the output logits being averaged and thresholded to gain the final segmentation. (**B**) Shows a sample of the raw images (left) along with the expert annotations for Rods and Rings used for training. It is worth noting that although these labels are provided by a highly skilled expert in the field, they are by no means an “absolute” ground truth; and contain some implicit biases and variations between the images.

Training was performed on a system with a NVIDIA GeForce RTX 3090 equipped using the Adam optimizer [16, 22] with an initial learning rate of 0.001 which decayed exponentially at a rate of 99.9% per epoch. During training, the loss for the training was recorded along with the loss on the testing dataset, although the latter was not used for hyperparameter optimisation to avoid overfitting and preserve integrity of the experiment.

##### Post-processing

After the models were trained, the masks were used to construct polygons for the rings and rods using a simple contour fitting algorithm [23]. For each image, several measurements were taken which were deemed biologically relevant (e.g. the number of rods/rings, the average perimeter and area of the rod/rings, the ratio of area to perimeter of rods and rings) to study the utility of the model in the biological context.

##### Testing

For testing the models on the original data, the final testing folds for each cross-validation set were predicted and compared against the ground truth binary masks provided by an expert annotator using both the Dice coefficient score and the per-pixel F1 score for pixel classification. In addition to the main dataset, we also tested the model on two further datasets: firstly, a time course dataset recorded with the same microscope as the primary dataset but in the context of an experiment examining the effect of differentiation media (N2B27) on the cell line. Here, the dataset is comprised of 306 images files at 11 different time-points including two control points. In this case, the density of rings and rods quickly diminishes over time **Fig. 3**. Testing the AI’s performance on this dataset not only tests its ability to segment images out of the training domain, but also serves as a simulation into how this model could be used in practice.

**Figure 3.**
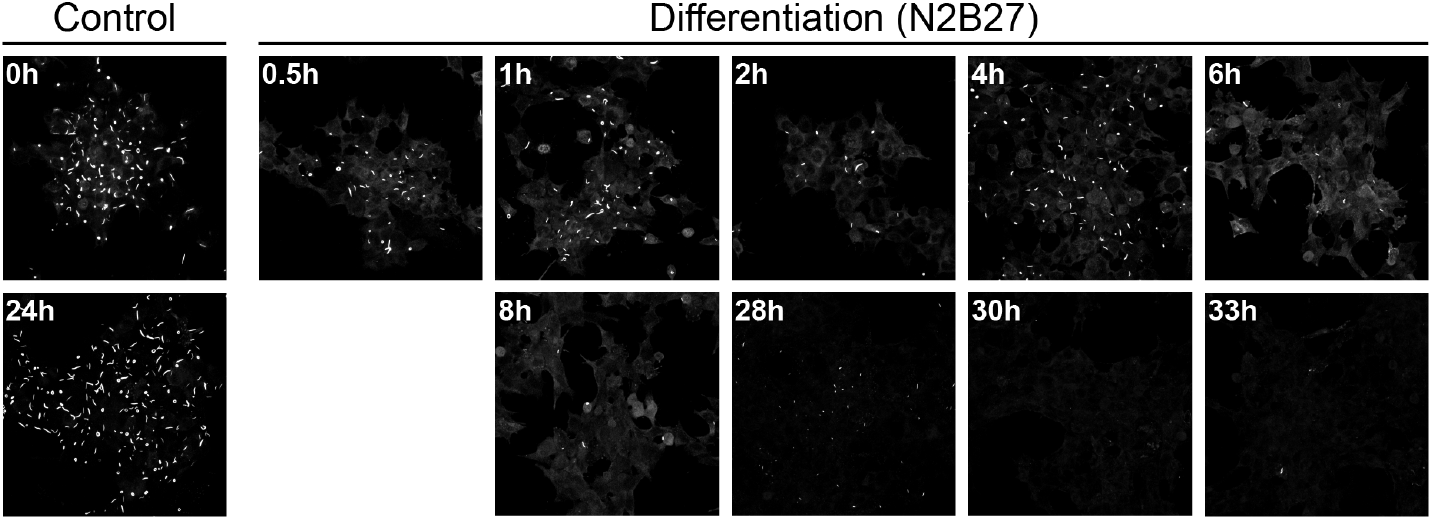
Differentiation of mouse ESCs and the loss of IMPDH2 RR structures. Examples of control cells at the start (0h) at 24h in pluripotency media (left), and cells over a period of 33h, sampled at the indicated time-points following transition into differentiation media (N2B27).

The final testing dataset examines the effect of a different microscope on the AI’s performance. In this case, we have 90 images recorded on a different microscope to what was used on the primary dataset. This dataset tests multiple out-of-domain factors including image scaling and different hardware image processing pipelines that are previously unseen in the primary dataset. For this set, two preprocessing pipelines were used for comparison, as the new microscope’s resolution was different to the original. The first, where the objective magnification was kept constant with the primary microscope, and the second, where the image was cropped so the resolution of the cells (i.e. the average pixel width) was kept constant to the original microscope **Fig. 4**.

**Figure 4.**
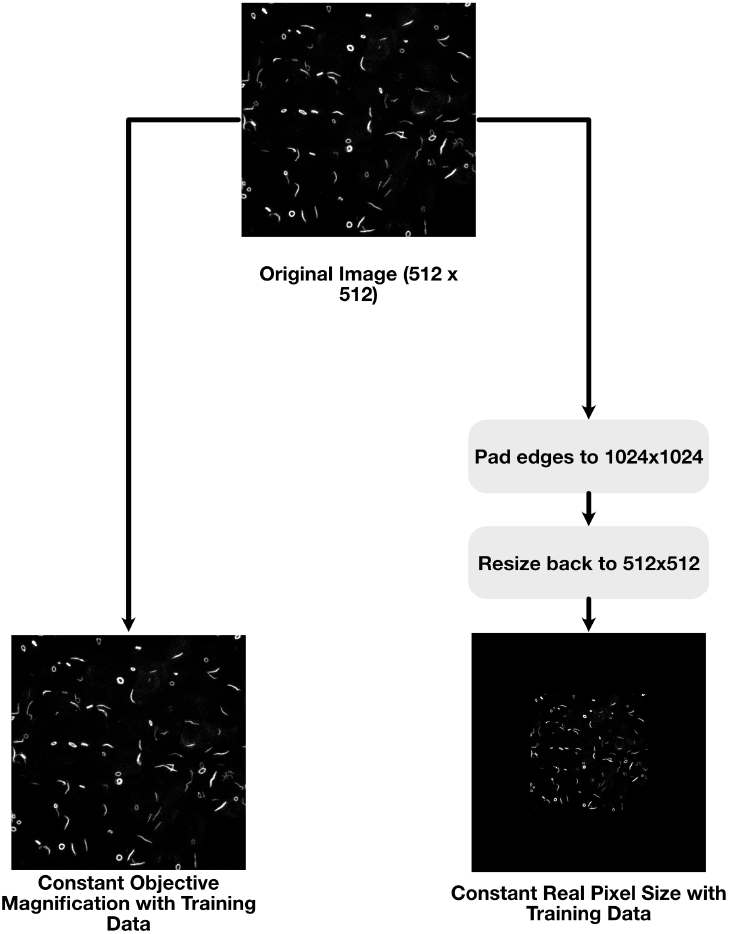
Preprocessing pipelines for datasets. In general, we found through development that very little preprocessing needed to be applied to the raw images before inputting into the model, with normalisation harming the segmentation results rather than helping. For the experiment with a change of microscope, two pipelines were developed due to a change in output resolution of the image: one fixing the objective magnification to that of the training data; and one fixing the real pixel size with that of the dataset. We find that the latter far outperforms the former.

For the time course and domain shift evaluations, since this data was not previously seen by the model; the ensemble model approach was used on the entire dataset, again recording the per-pixel F1 score and Dice coefficients per image. In both cases, the logit maps were also used to construct receiver operating characteristic (ROC) curves for each image, then averaged over the image set, giving a mean ROC curve along with a standard deviation range. Finally, for each model output, the summary metrics after post-processing were compared with those calculated from the ground truth; to test for overall utility of the model in a biological context. This was achieved by calculating the *R*^2^ score for each measurement to test for the strength of correlation of the measurements from the ground truth to the model output.

##### Image Processing Pipeline

Images from the Andor Dragonfly Microscope were exported as .ims files, and maximum intensity projections were created (1024 × 1024 pixels) before being converted to .jpgs for rod ring prediction to be made off from the model. For instances in which annotations were required, OpenCV [24] was used to draw initial contours off single channel maximum projection images. These were then manually cleaned and separated into rod and ring label layers in napari [25] which were exported as binary masks for training.

### Biological Methods

#### Routine mouse embryonic stem cell culture and differentiation

Wild-type E14-Tg2A mESCs were seeded at a density of 1.2×104 cells/cm2 on 0.1% (v/v) gelatin-coated flasks in ESL medium which is comprised of GMEM (Gibco) supplemented with 15% Foetal Bovine Serum (Gibco Cat. No. 10270-106), Non-Essential Amino Acids (Gibco), Sodium Pyruvate (Gibco), Glutamax™ (ThermoFisher, Cat. No. 35050038), β-mercaptoethanol (Gibco, Cat. No. 31350010; final concentration 0.05 mM), and LIF (QKine; QK018) in a humidified incubator maintained at 37°C with 5% CO_2_. Cells were passaged every other day using TrypleE as a dissociation reagent. The culture medium was changed on non-passage days. For differentiation assays, cells were seeded directly in N2B27 [26, 27] and medium changed each day [28]. N2B27 was kept at 4°C protected from light and was used within three weeks. Cells were tested monthly and certified negative for mycoplasma.

#### Immunofluorescence

Cells were seeded at a density of 3.0×104 cells/cm2 in 8-chambered Ibidi slides in ESL for 24h [26, 27]. For samples that were gathered for the differentiation time course experiment, ESL media was exchanged for N2B27 at 24 hours and cells were fixed at the defined intervals. Samples were then processed for immunofluorescence as previously described [10]. Prior to fixing cells were washed with phosphate buffered saline without Calcium and Magnesium (PBS^-/-^). Single time point experiments were fixed for 30 mint at room temperature with 4% formaldehyde (Thermo Scientific/Pierce). Samples part of the differentiation time course were fixed at the defined intervals with 4% formaldehyde for 10 minutes at 37°C. After fixation, cells washed multiple times (10 min each) with PBS^-/-^-supplemented with 10% FBS and 0.2% Triton X100 (PBSFT), followed by three hour-long washes in PBSFT to block and permeabilise the cells. Cells were then incubated with an antibody raised against IMPDH2 (12948-1-AP, Proteintech) diluted 1:2000 in PBSFT at 4°C over-night. Cells were then rinsed three times with PBSFT and exposed to three lots of one-hour washes in PBSFT before incubation over night with alexa-488 conjugated secondary antibody diluted 1:500 in PBSFT with Hoechst (1:1000) to mark nuclei at 4°C. Cells were then rinsed with PBS supplemented with 0.2% FBS and 0.2% Triton X100 (PBT) three times before three hour-long PBT incubations at room temperature. After the final rinse, cells were mounted in ProLong Antifade (P36980, Thermo Fisher Scientific) and stored at 4°C until imaging.

#### Confocal Microscopy

Fixed and stained cells were primarily imaged on an Andor Dragonfly spinning disc confocal microscope mounted on an inverted Leica DMi8 base using a 40x 1.4 NA oil-immersion objective. Hoechst and Alexa-647 were sequentially excited with 405 and 637nm laser diodes respectively, and emitted light reflected through 450/50 nm and 700/25nm bandpass filters respectively. An iXon 888 EM-CCD camera was used to collect emitted light, and data were captured using Fusion version 5.5 (Bitplane). For microscope comparison tests, samples were imaged on a Zeiss LSM800 Airyscan confocal microscope mounted on an inverted Axio Observer Z1 base using a Plan-Apochromat 63x 1.40NA Oil-immersion objective. Alexa-647 was excited with 640nm laser diode, with emitted light reflected through a variable secondary dichroic beamsplitter set between 656-700nm. Emitted light was directed to a GaAsP photomultiplier tube, and data captured using Zen Blue version 2.6.

## Results

### Primary dataset

The model training progress shows convergence of the models on the training datasets while still not overfitting **Fig. 5A**. After training, the models were tested on each testing fold using the Dice and F1 scores (**Fig. 5B**), and the logit maps were used to construct ROC curves **Fig. 5D**. In both cases, we see high performance from the model, achieving an overall average Dice scores of 0.806±0.054 and 0.809±0.035 for rods and rings respectively and F1 scores of 0.806±0.054 and 0.809±0.036 for rods and rings respectively. We also see a strong performance in the ROC curve analysis, achieving a mean AUC of 0.8965 and 0.9063 for rods and rings (**Fig. 5D**). In representative examples of model outputs (**Fig. 5C**), the model clearly segments the images well, apart from in a few select areas of disagreement that may be a result of annotation bias (the correlation of derived measurements in are also shown (**Fig. 5E,F**; and **Supplemental Fig. S1**). Overall, we see a strong correlation between the researcher derived measurements and those produced by the model for the number of rods and rings (*R*^2^ of 0.7255 and 0.8572), but much lower for the other measurements (**Supplemental Fig. S1**), showing that the output of the AI can be used to track some biologically meaningful information within the domain of the dataset.

**Figure 5.**
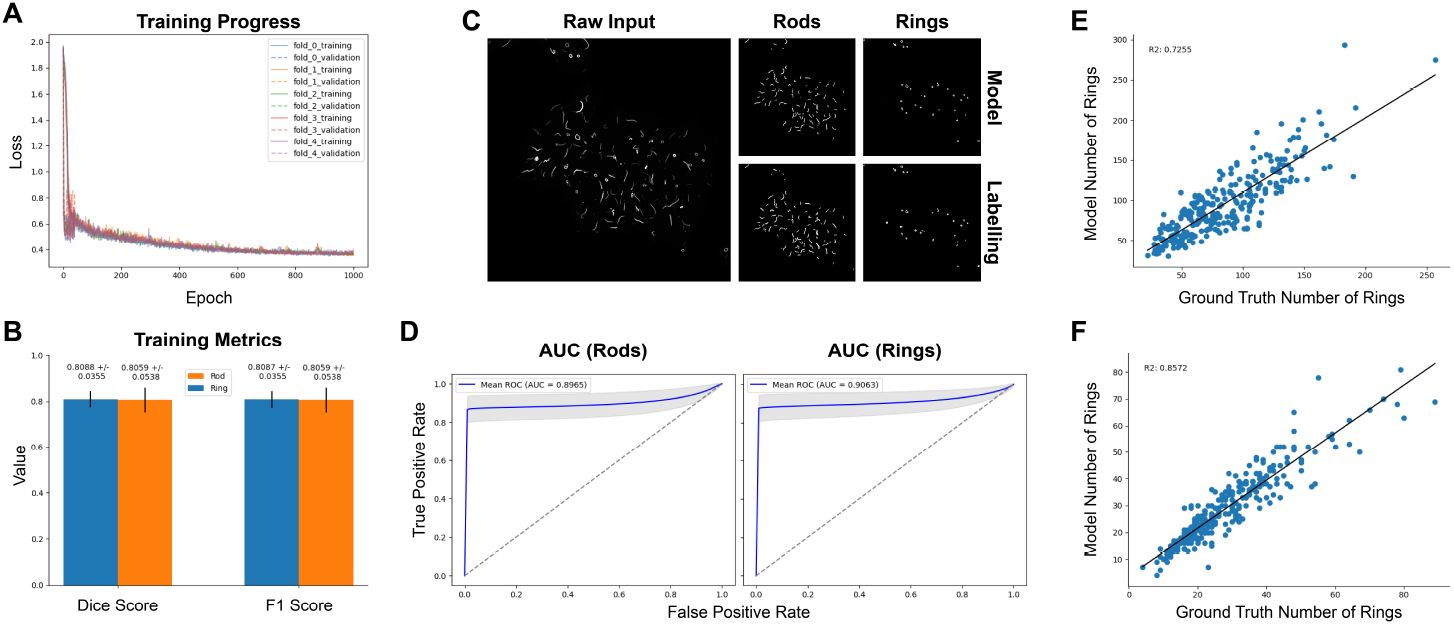
Model Training. (**A**) shows the training and testing losses of each of the models over time. We see that the models similarly converged over time, showing there were no training folds significantly different to others for training. (**B**) shows the mean Dice and F1 scores for the models tested on the holdout test set for the rod and ring mask recovery. (**C**) Shows a representative example outputs along with the relative labels for the rods and ring masks. (**D**) Shows the per-pixel AUC curve for each of the mask. (**E**) and (**F**) show the correlation of the derived measurements from the model segmentations versus the segmentation provided by an expert annotator. A strong correlation here shows not only an accurate segmentation, but also a biologically helpful model that can be used for downstream analysis.

### Time Course Dataset

To test the model’s performance at predicting data outside of the training domain, an additional dataset was recorded that examined the effect of differentiation media on the rid-ring structures over time, which permitted us to test the out of domain performance of the model. In this case, we again record the Dice and F1 scores (**Fig. 6A**) and show similar performance to the testing dataset albeit with some reduction in images with low numbers of rods and rings, with average dice scores of 0.7189±0.0659 and 0.7338±0.0484 for rods and rings respectively and F1 scores of 0.7186±0.0660 and 0.7334±0.0484 for rods and rings respectively. ROC curve analysis shows similarly high AUC scores for rods and rings compared to the primary dataset, with AUCs of 0.734 and 0.856 for rods and rings (**Fig. 6B**).

**Figure 6.**
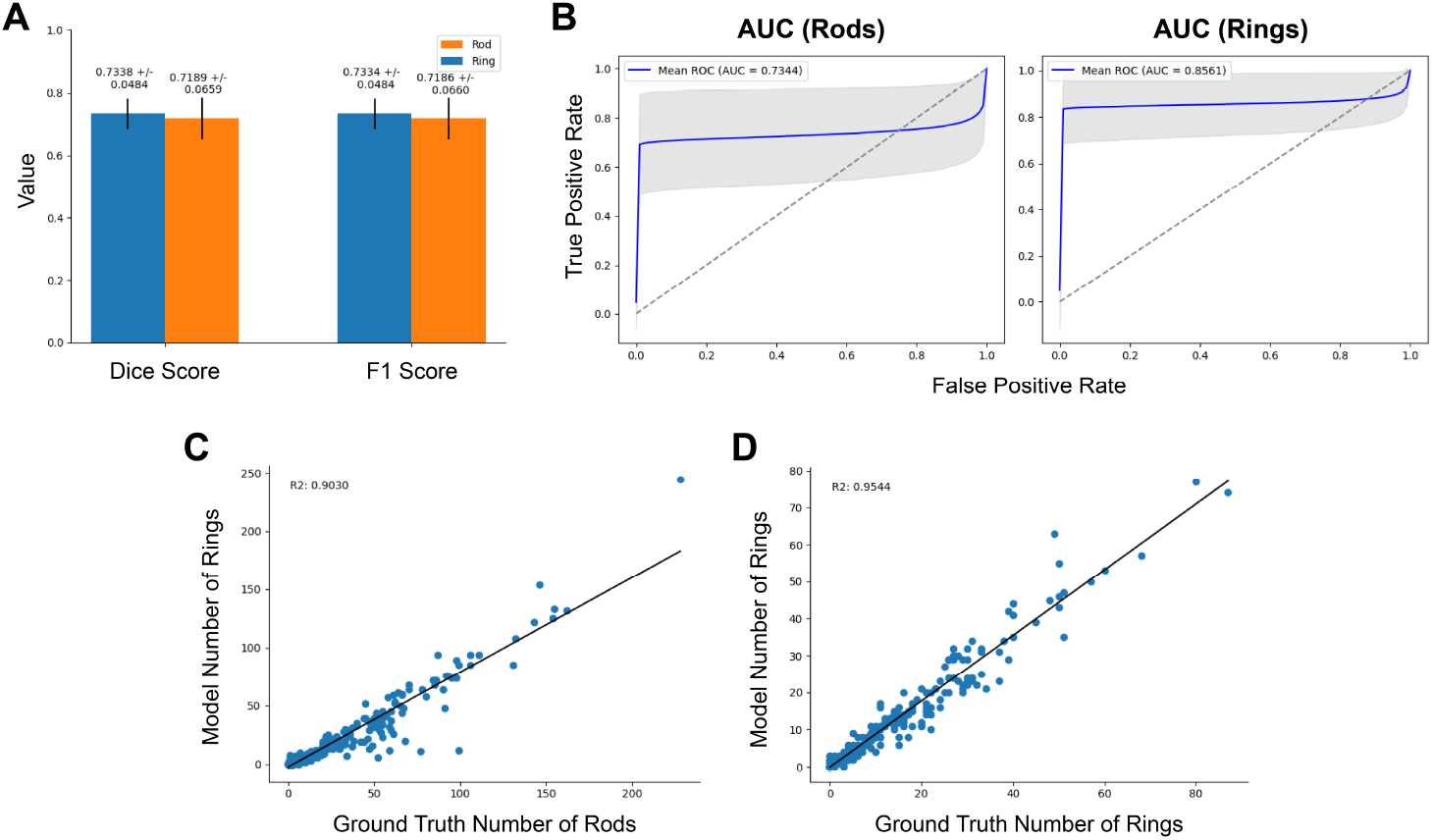
Time-course metrics. For the time-course dataset, the model trained on the primary dataset was applied via an ensemble mechanism with the resulting metrics to test for out-of-domain performance. (**A**) shows the Dice and F1 scores for rods and rings for each image compared with expert annotation. (**B**) shows the per-pixel AUC scores for both the rods and rings. (**C**) and (**D**) show the correlation between derived features from the segmentations for both the expert analysis and the model. As with the previous primary dataset, correlation here shows the model can recover meaningful biological information as well as pixel-wise accuracy.

There is a high correlation between the ground truth expert-annotated masks and the output of the ensemble model, with *R*^2^ of 0.9030 and 0.9544 for rod and ring counts respectively (**Fig. 6C,D**). This indicates that the model is robust to changes in rod ring morphology due to biological changes in the experimental conditions. The plots of several measurements split by their time point can be seen in (**Fig. S2**). We show that the model is successfully able to find similar patterns in the time course experiment as the human annotations.

### Microscopy Change Dataset

The model was tested on a dataset recorded by a different microscope to examine the robustness of the model to changes in the nature of the images. These data were pre-processed in two ways as the microscope resolutions differ: one pipeline to keep the magnification constant with the primary dataset and another to keep the pixel height and width constant (**Fig. 4**). When examining the performance of the model, we see relatively low Dice scores of 0.2795 ± 0.101 and 0.3483 ± 0.127 and F1 scores of 0.3480 ± 0.127 and 0.2798 ± 0.101 for rods and rings respectively (**Fig. 7A,B**). Additionally, ROC analysis for the constant magnification dataset shows similarly disappointing results (**Fig. 7B**), with AUCs of 0.5047 and 0.5471 for rods and rings respectively. Representative examples qualitatively show that the model has unsuccessfully read these images as there are significant artefacts with the output masks (**Fig. S3**).

**Figure 7.**
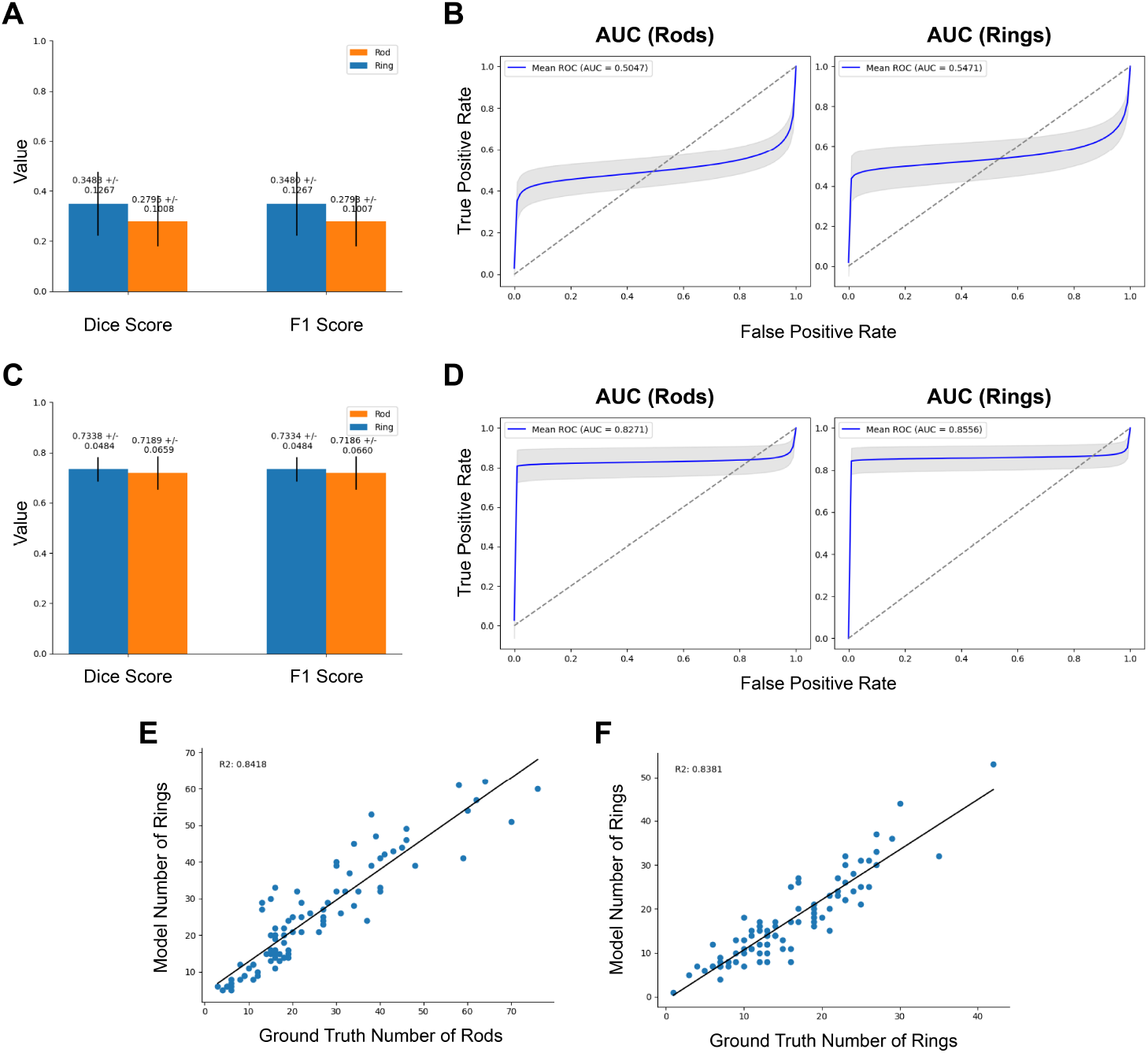
Microscopy experiment metrics. Data from a different microscope was used to test for application of the model out-of-domain. We find that when preprocessing fixes the objective magnification of the input data, the model performance is fairly poor. The Dice and F1 scores for rod and ring segmentation compared to expert analysis are shown (A), as well as the pixel-wise AUCs for rods and rings (B). By preprocessing the data to preserve the real pixel size compared to the primary dataset, we see much better model performance. (C) shows the Dice and F1 scores for the rod and ring segmentations after this preprocessing step and (D) shows the AUC scores. (E) and (F) show the correlation of the latter preprocessing pipeline model with the expert annotations. We again see a strong correlation with the expert annotation showing strong biological relevance of the model.

Interestingly, for the pixel height and width constant preprocessing pipeline, we see much better results (**Fig. 7C**), with higher Dice scores of 0.719±0.066 and 0.733±0.048 and F1 scores of 0.718±0.066 and 0.733±0.048 for rods and rings respectively. Furthermore, in this case the ROC analysis shows much greater performance (**Fig. 7D**), with AUCs of 0.8271 and 0.8556 for rods and rings. Lastly, for the pixel height and width constant preprocessing pipelines, we also measure the correlation between expert-annotated mask measurements and the AI model output mask measurements (**Fig. 7E,F**). We see a strong correlation between the two, with an *R*^2^ score of 0.8418 and 0.8381 for rod and rings counts.

## Discussion

In this work, we have developed a fully automatic end-to-end image segmentation pipeline for rod ring microscopy images that can derive biologically relevant measurements over the course of an experiment. However, we still feel like there are several open questions for this work:

### Comparison to Existing Models

Pre-trained transformer-based models such as Segment Anything (SAM) [17], You Only Look Once (YOLO) [16] and Segformer [10] are all state-of-the-art models in their respective fields, however all have significant drawbacks in this domain. SAM still requires some user input in detecting objects for segmentation; so, inclusion of the model alone in the pipeline would lose the automatic nature of the work. Since the throughput of microscopy can be high, this is crucial to the utility of the model. Furthermore, **Fig. 8** shows examples of images segmented with SAM showing significant errors in its segmentations. Cellpose is a similar solution, designed to be general to cellular applications, however we find similar issues with the model’s ability to segment rods and rings (**Fig. 8**). Similar issues exist for YOLO and Segformer. Both of these models segment without user input, but use a pre-defined set of labels that do not include rods and rings. In development, retraining or finetuning these models was unsuccessful due to overfitting, perhaps due to the large size of these models compared with the relatively small size of our dataset.

**Figure 8.**
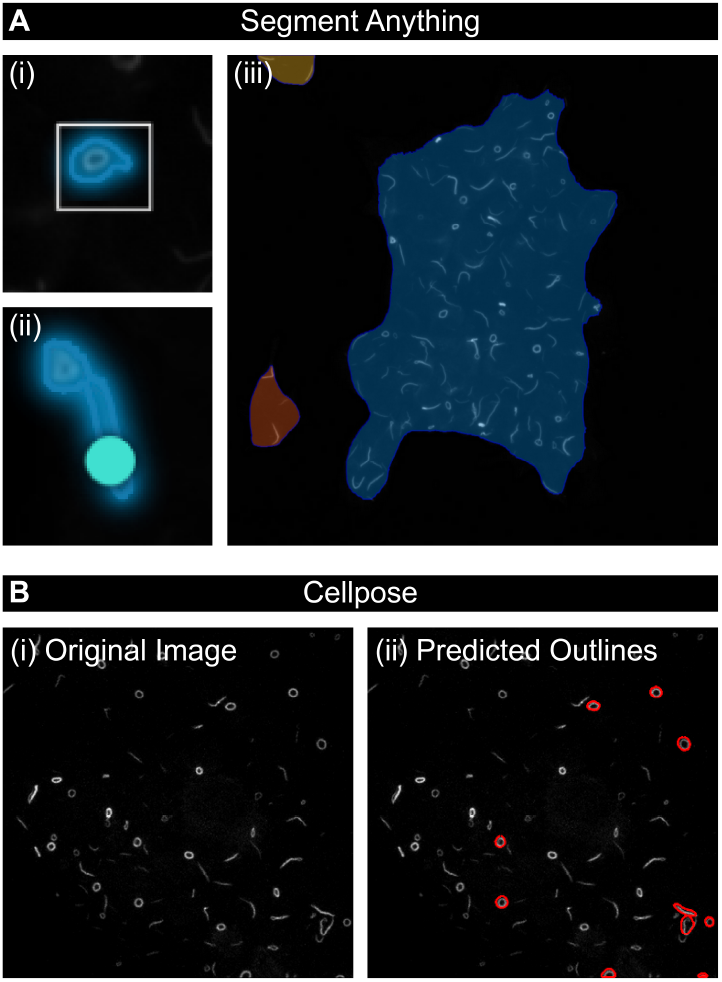
Example Segmentations from Segment Anything and Cellpose. (**A**) Segment anything provides context-free segmentations using user prompts to drive the segmentation process, however we find that using the different modes for SAM we get poor results for our specific datasets. (i) shows the segmentation using the box-prompt mode and (ii) show the segmentation using the point prompt mode. The representative example (ii) showing the model unable to differentiate between a rod and a ring object. (iii) shows the “Whole picture” approach which fails to single out individual objects in the image. (**B**) The original image (i) and predicted outcomes (ii) using Cellpose showing an inability to segment rods and rings correctly.

### Measurement Variance

Measuring correlation with expert annotations is dangerous for several reasons: firstly, it assumes that the expert annotation is a completely accurate source of truth (see below). Secondly, it is also (with the methods discussed in this work) weak to high variance measurements: for example, due to the resolution of the data, large changes in some measurements can occur when only small changes are made in the annotation. In this work, the model showed strong correlation between expert annotation and model output for ring and rod counts. This measurement shows lower variance when the sample size (i.e. the number of rings and rods available) is high. Since the microscope has a fixed resolution, this occurs when there are a large number of small, low-pixel count rings and rods available. Unfortunately, measuring the area or perimeter of the rings and rods is difficult in this case.

When the rods and rings have a small number of pixels, an additional pixel labelled by the model results in a large change in the area and perimeter measurements. Additionally, the low resolution of these objects results in only a few possible values; leading to discontinuities in histograms when performing downstream analysis (**Fig. 9**). The solution to the above problem is to take images where fewer rings and rods are visible, but at a much higher resolution, where the area and perimeters can be measured with a higher precision for analysis. However, this reduces the number of rings and rods visible, which itself increases variance for these measurements given the same number of images taken.

**Figure 9.**
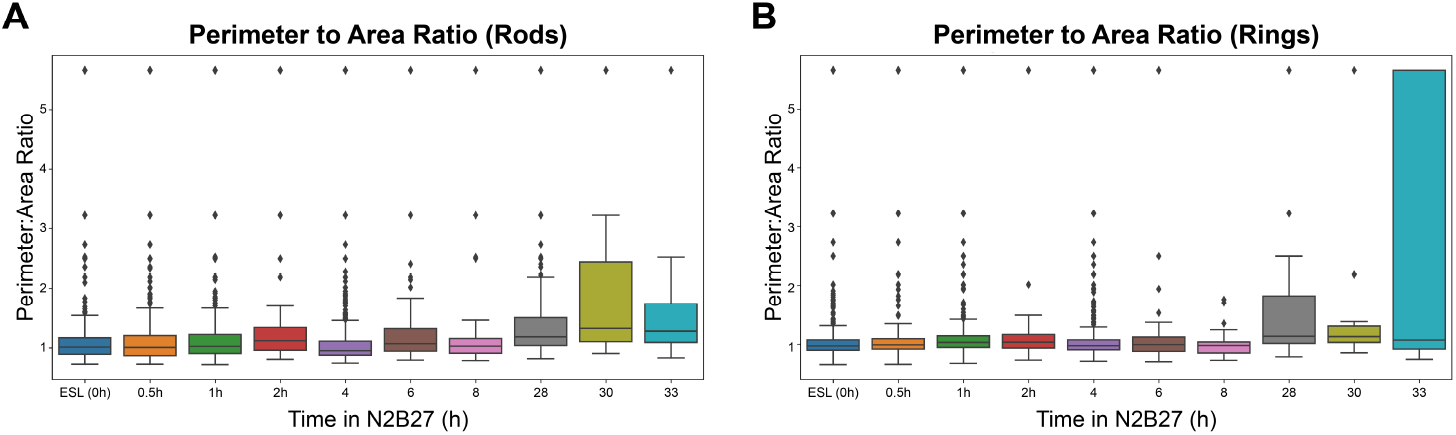
Perimeter to Area Ratio for the Time Course Experiment. For each of the detected polygons, the perimeter and area were calculated, to account for changes due to the change in media. However, due to the pixel size of the objects in the microscopy images, we see common ratios appear due to noisy segmentation. In future work, if this ratio is to be used, a higher resolution is needed for each of the objects to allow for more granularity in the perimeter and area calculations.

### Expert Bias

In the recording of metrics for this work, we encounter a common problem in using AI for biological or medical tasks which was the lack of an objective ground truth. While the results in this work are promising, these are metrics based on a comparison to expert annotation that may include biases and variances between and within the datasets that make annotations inconsistent. Some objects close to the edge of the image or objects slightly dimmed compared with other objects, might be labelled as a ring or a rod by the expert but not by the AI model; or vice versa. This leads to some slight inconsistencies with the metrics, and it is arguable as to whether areas the model was marked as incorrect were incorrect at all (**Fig. 10**).

**Figure 10.**
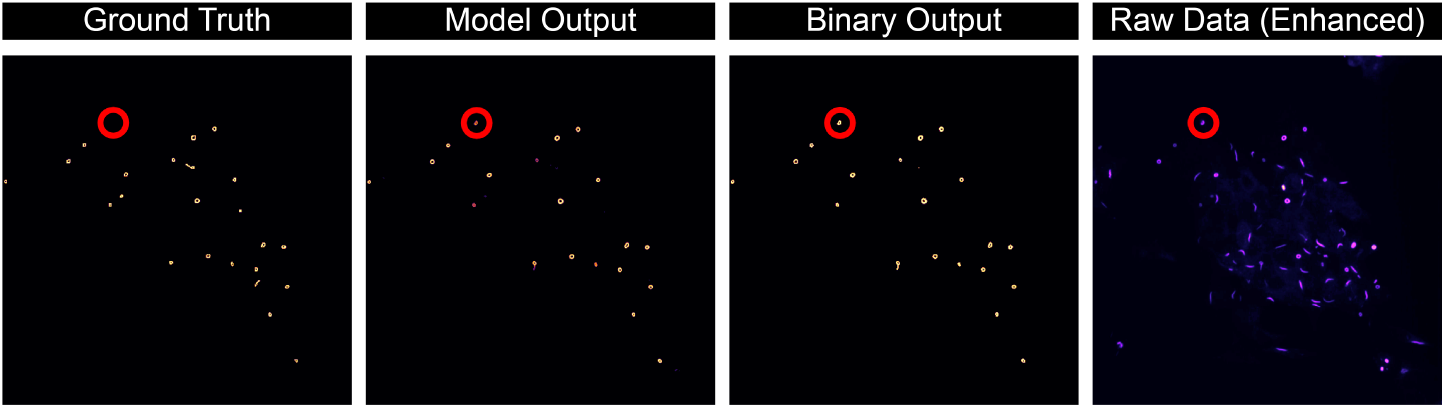
Comparison of model labelling rings potentially missed by experts. Examples of ground truth labelling done by out microscopy expert, the model logit “fuzzy” output mask, the binary mask for the images, and the original image with increased exposure. We see that a ring-like structure visible in the original image (in red) is picked up by the model but not annotated by the expert annotator.

### Microscope Resolution

In testing for the effect of changing the microscope on the AI model performance, we see that the effective method to preserve model performance is to preprocess the data in a bespoke way that preserves the real-life height and width of a pixel in space. For other microscopes, this can be calculated easily but is crucial for model inference as shown above. Furthermore, we see a trade-off within the measurements we take, and the resolution of the images taken from microscopy: if the microscope is at a lower magnification, the resulting image will contain a high number of objects, but at low resolution. This results in a lower variance in the “object-aggregate” measurements such as the count of rods or rings, but a lower accuracy in the “within-object” measurements such as area or perimeter. This is because at a low resolution, each object may only take a small number of pixels in the image, and therefore the resulting area may be under 5 pixels large.

On the other hand, if the magnification is significantly high (and the model retrained to account for this) – we would see much more accurate reporting of measurements such as area and perimeter; however, the number of objects identified would be far fewer; increasing variance in this way.

### WebApp

Crucial to the development of this model was the development of an accompanying webapp, allowing feedback from expert biologist in how to improve the model in the biological context. **Supplemental Fig. S4** shows a screenshot of this model in practice, and we have made the webapp code available for use by research teams in an easy to deploy manner.

## Conclusion

We have developed an AI tool with an accompanying webapp to identify, classify, and quantitatively analyse IMPDH2 RR structures from confocal microscopy images. Furthermore, we’ve shown that our model works out of domain when images are taken on different microscopes, and when the size and conformation of RR structures differ (e.g. during differentiation).

The RR conformation is not unique to IMPDH2, nor are these structures limited to mammalian cells: depending on the biological context, different metabolic enzymes can form these structures which can vary in size, frequency, and topology. Examples include CTPS in yeast [8, 29], IMPDH2 in zebrafish teeth [30], and various filamentous RR proteins in drosophila and yeast [8]. Additionally, other RR-like structures, distinct from both IMPDH2 RRs and CTPS1 cytoophidia have been identified such as nematosomes [31, 32] and loukoumasomes [15, 33], the latter enriched cytoskeletal proteins. Therefore, the pipeline we have developed can be of broader significance to multiple groups, allowing them to train their own datasets and quantitatively analyse these structures.

Finally, biological systems do not exist in static form, and many proteins are highly dynamic. Indeed, temporal imaging of fluorescent CTPS1 and IMPDH2 has shown that RRs are dynamic structures, coalescing into rods and rings, and increasing/decreasing their size over time [7, 34]. Our model at present does not cater for time-lapse tracking of live RRs in real-time, but as segmentation and classification is usually the bottleneck, tracking segmented RRs could be a feasible modification for future pipeline developments.

## List of Abbreviations

AI: Artificial Intelligence
CTPS1: Cytidine Triphosphate Synthase 1
ESC: Embryonic stem cell
FBS: Foetal Bovine Serum
GMP/GTP: Guanine monophosphate/Guanine Triphosphate
IMPDH2: Inositol monophosphate dehydrogenase 2
PBS^-/-^: Phosphate buffered saline without calcium or magnesium ions
PBSFT: PBS supplemented with 10% FBS and 0.2% Triton X100
PBT: PBS supplemented with 0.2% FBS and 0.2% Triton X100
RR: Rod and Ring

## Acknowledgement

We would like to thank Anna Bigas for the kind gift of the E14-Tg2A mouse ESCs used in this study. We are indebted to the University of Liverpool’s (UoL) Centre for Cell Imaging (CCI) facility for provision of state of the art imaging equipment funded through grants awarded by the MRC (MR/K015931/1) and the BBSRC (BB/M012441/1, BB/R01390X/1). We would also like to thank Thomas Waring from the CCI for excellent technical support, assistance, and training. Additional thanks are extended to members of the DAT lab for useful discussions and support.

DAT was funded in this work by the BBSRC (a DTP PhD studentship BB/T008695/1, project reference 2599454 supporting MJH, and a New Investigator Grant BB/X000907/1), the NC3Rs (NC/P001467/1), The Royal Society (RGS/R2/202075), a Wellcome Trust non-clinical ISSF, and UoL’s Technology Directorate voucher scheme. KFJ was supported in this work by core funding from Department of Electrical and Electronic Engineering for YT. SB and YZ were funded by the Wellcome Trust’s Investigator Award in Science Programme. Grant Number: 222530/Z/21/Z. None of the funding bodies had any input in the design of the study or data collection, analysis, and interpretation, or in writing the manuscript.

## Author Contributions

STMB conceived the AI approach, developed, trained, and tested the software, analysed microscopy images, and co-wrote the first draft of the manuscript with MJH. MJH performed the experimental work and the associated microscopy, analysed the data, and annotated all microscopy images for training the model. KFH conceived the preliminary AI study and with DAT, and supervised YT who contributed to an initial feasibility study as part of his 3rd year research project. MRHW, NDP, and DJW had intellectual input into the underlying biological study and provided secondary supervision to MJH. YZ provided significant intellectual input into the AI methodology and provided comments on the manuscript. DAT conceived and obtained funding for the experimental study, led and managed the project, and wrote the manuscript. All authors read and approved the final manuscript.

## Data Availability Statement

All software, and code generated or analysed during this study are included in this published article’s supplementary information files and have been deposited in github https://github.com/gastruloids/gandalf. Test images use to train the data are also included.

## Supplementary Material

**Supplemental Figure S1.**
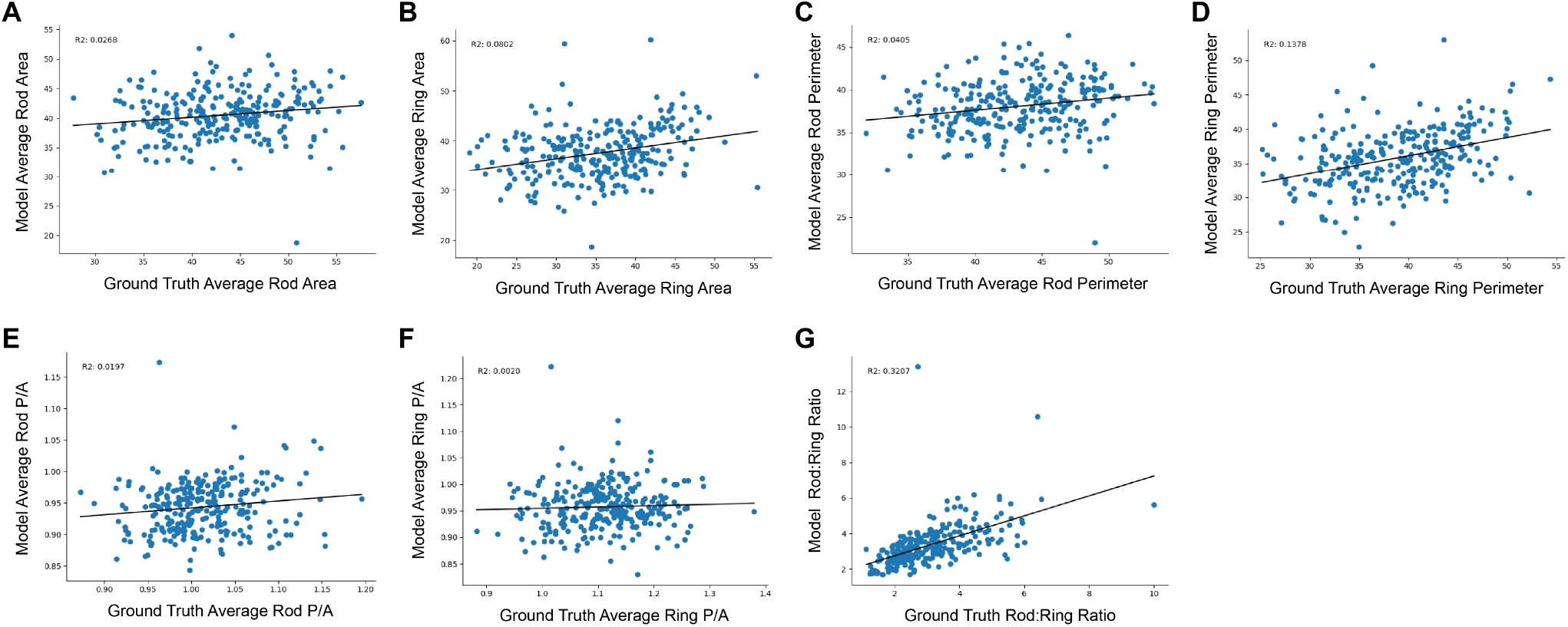
Additional Metrics for model training. Each graph shows ground truth measurements on the x-axis vs the model’s (y-axis) ability to capture the average area (A,B), and average perimeter (C,D), perimeter/area (P/A; E,F) for either rods or rings. The Rod:Ring ratio is also shown (G). *R*^2^ values are shown in each graph showing the correlation between the model’s ability to capture ground truth measurements.

**Supplemental Figure S2.**
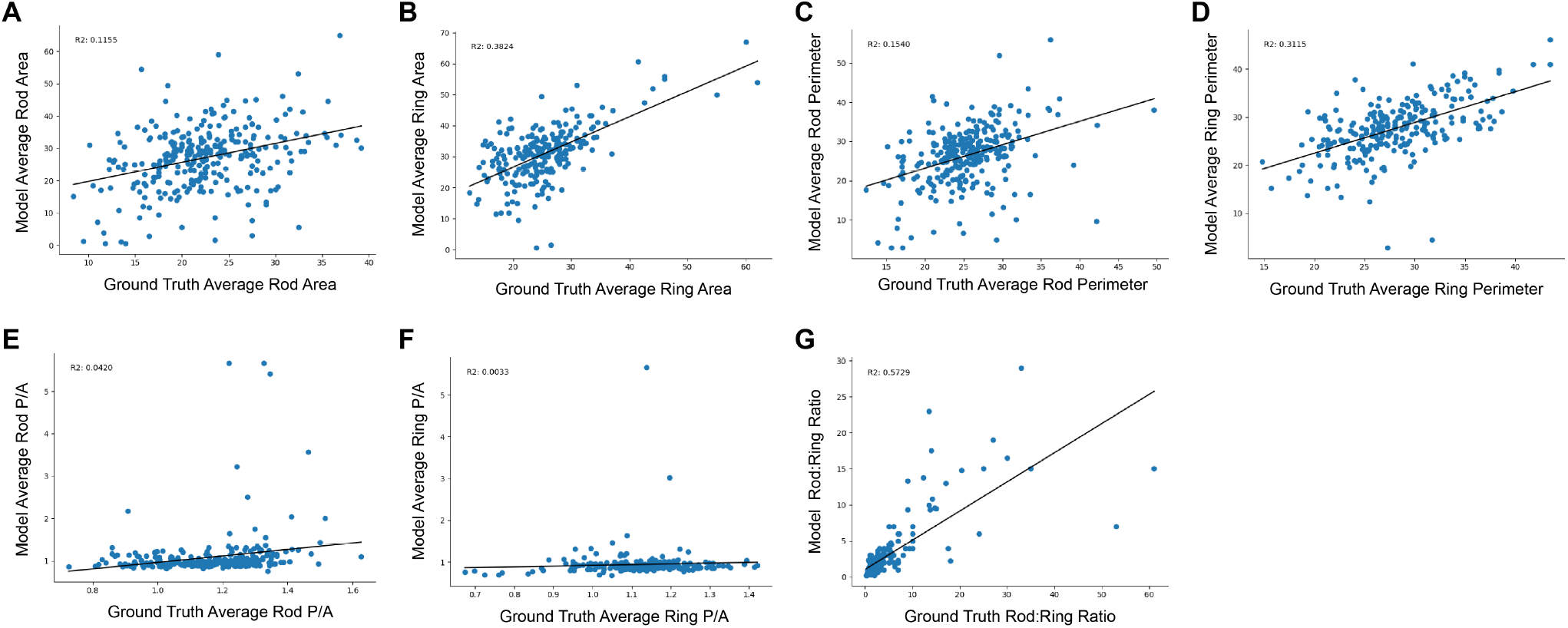
Additional Metrics for the time-course. Each graph shows ground truth measurements on the x-axis vs the model’s (y-axis) ability to capture the average area (**A,B**), and average perimeter (**C,D**), perimeter/area (P/A; **E,F**) for either rods or rings. The Rod:Ring ratio is also shown (**G**). *R*^2^ values are shown in each graph showing the correlation between the model’s ability to capture ground truth measurements.

**Supplemental Figure S3.**
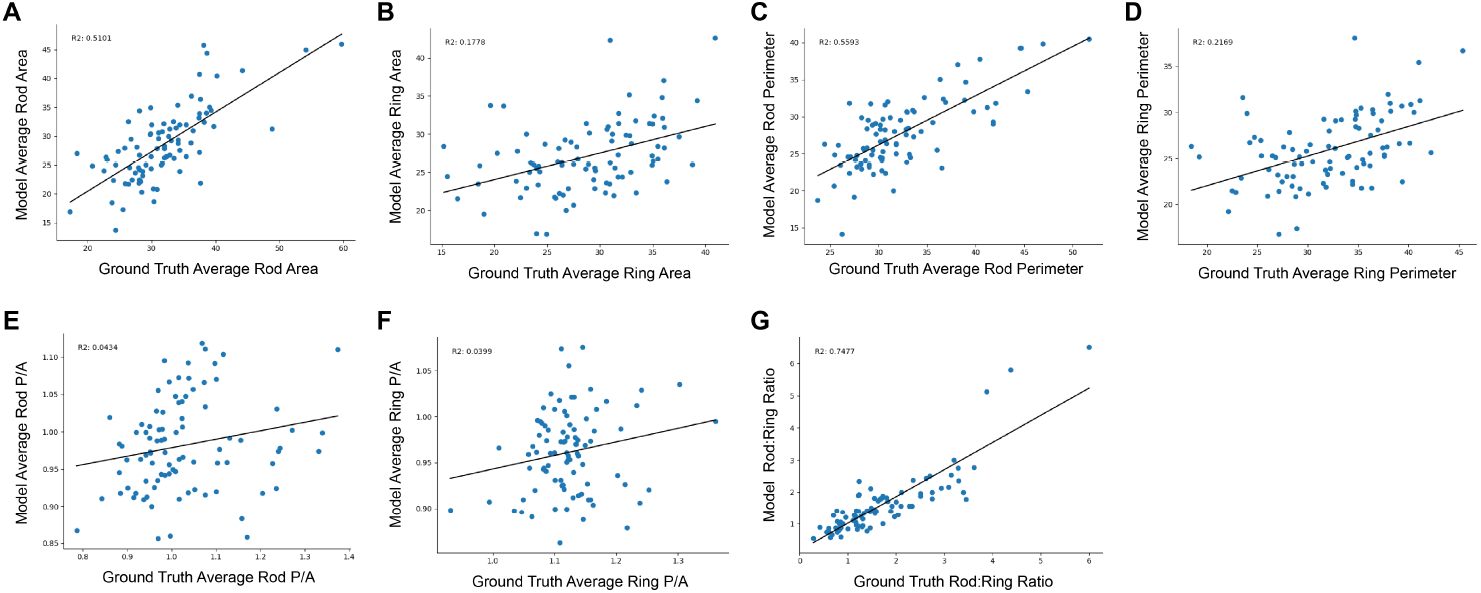
Additional metrics following modification of the microscopy pipeline. Each graph shows ground truth measurements on the x-axis vs the model’s (y-axis) ability to capture the average area (**A,B**), and average perimeter (**C,D**), perimeter/area (P/A; E,F) for either rods or rings. The Rod:Ring ratio is also shown (**G**). *R*^2^ values are shown in each graph showing the correlation between the model’s ability to capture ground truth measurements.

**Supplemental Figure S4.**
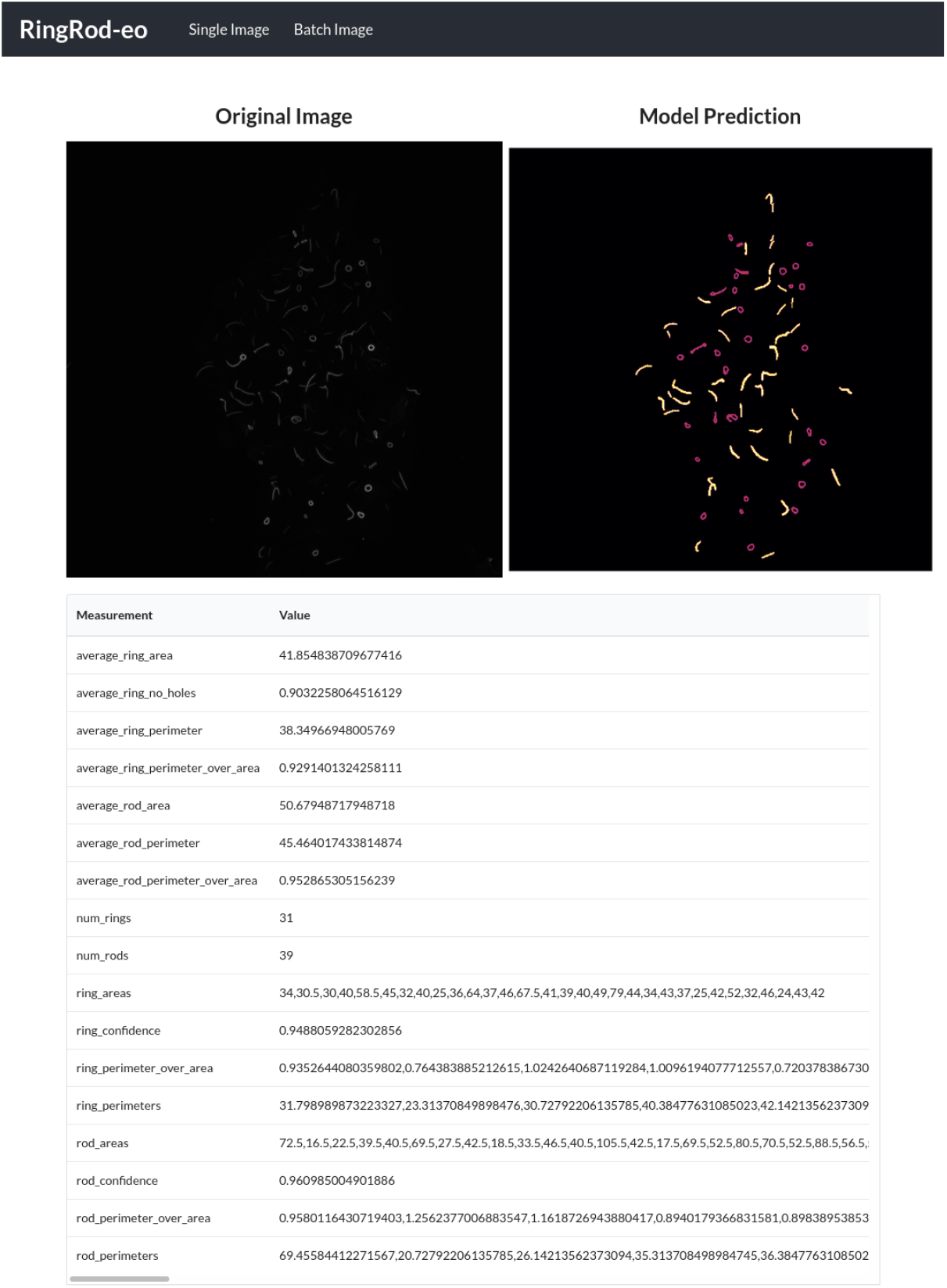
Webapp developed for lab use. During this work, constant feedback was needed from microscopists for successful development of the mode. This WebApp was developed for use by microscopists to automatically segment the confocal microscopy image files and calculate the summary features from the segmentation. Code for self-hosting the web app, along for the code used to develop the models can be found here: https://github.com/gastruloids/gandalf.

